## Supplemental Figures for "Dissecting the Human Leptomeninges at single-cell resolution"

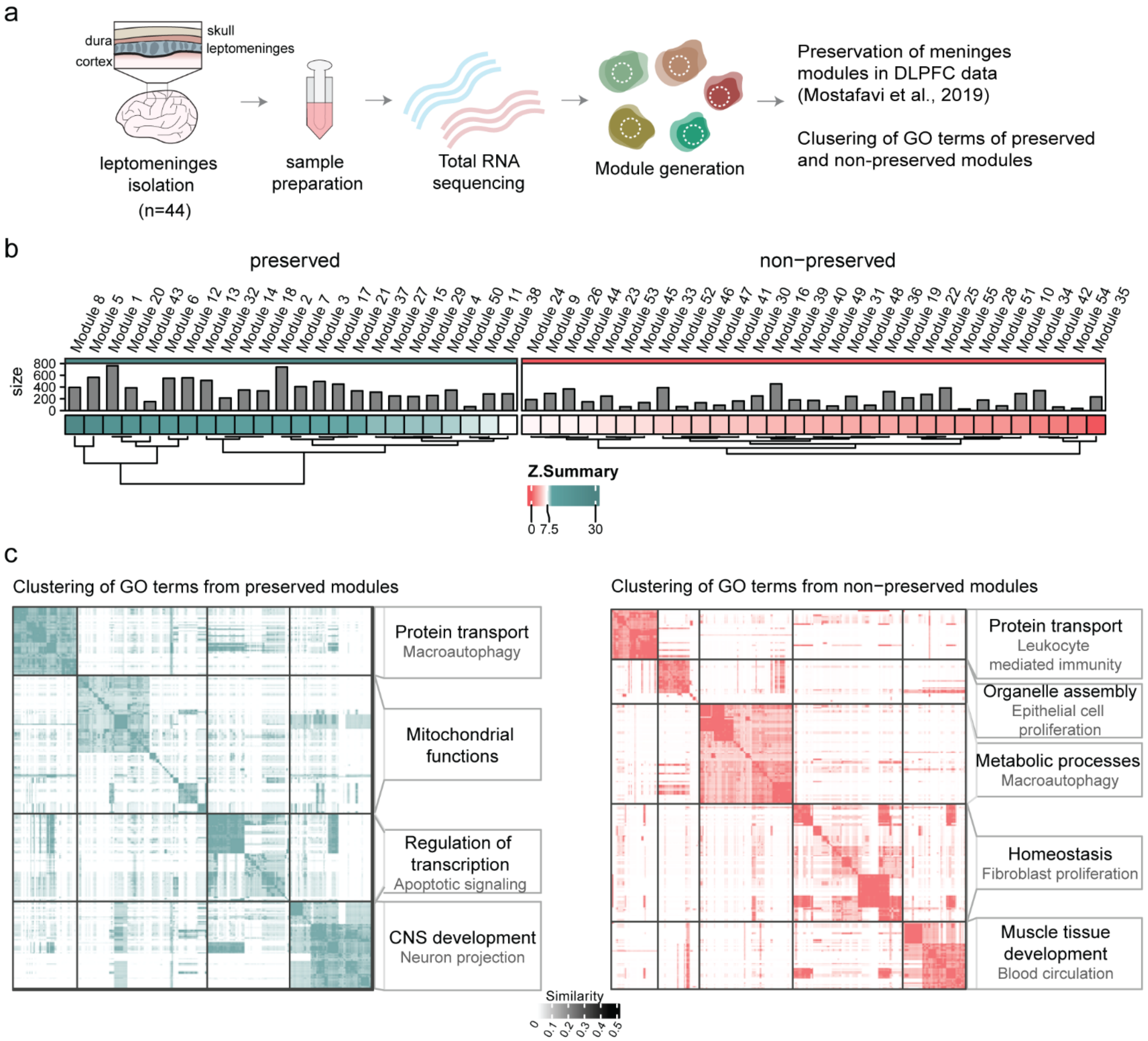


**Extended Data Figure 1. Gene co-expression modules indicate preserved and non-preserved functions of the human leptomeninges.** (a) Schematic illustration of experimental design and bulk RNA-seq analysis. (b) Co-expressed gene modules generated using SpeakEasy, and their gene set size and preservation scores in the DLPFC data from Mostafavi et al., 2019. (c) Clustering of gene ontology terms from preserved and non-preserved leptomeninges modules.

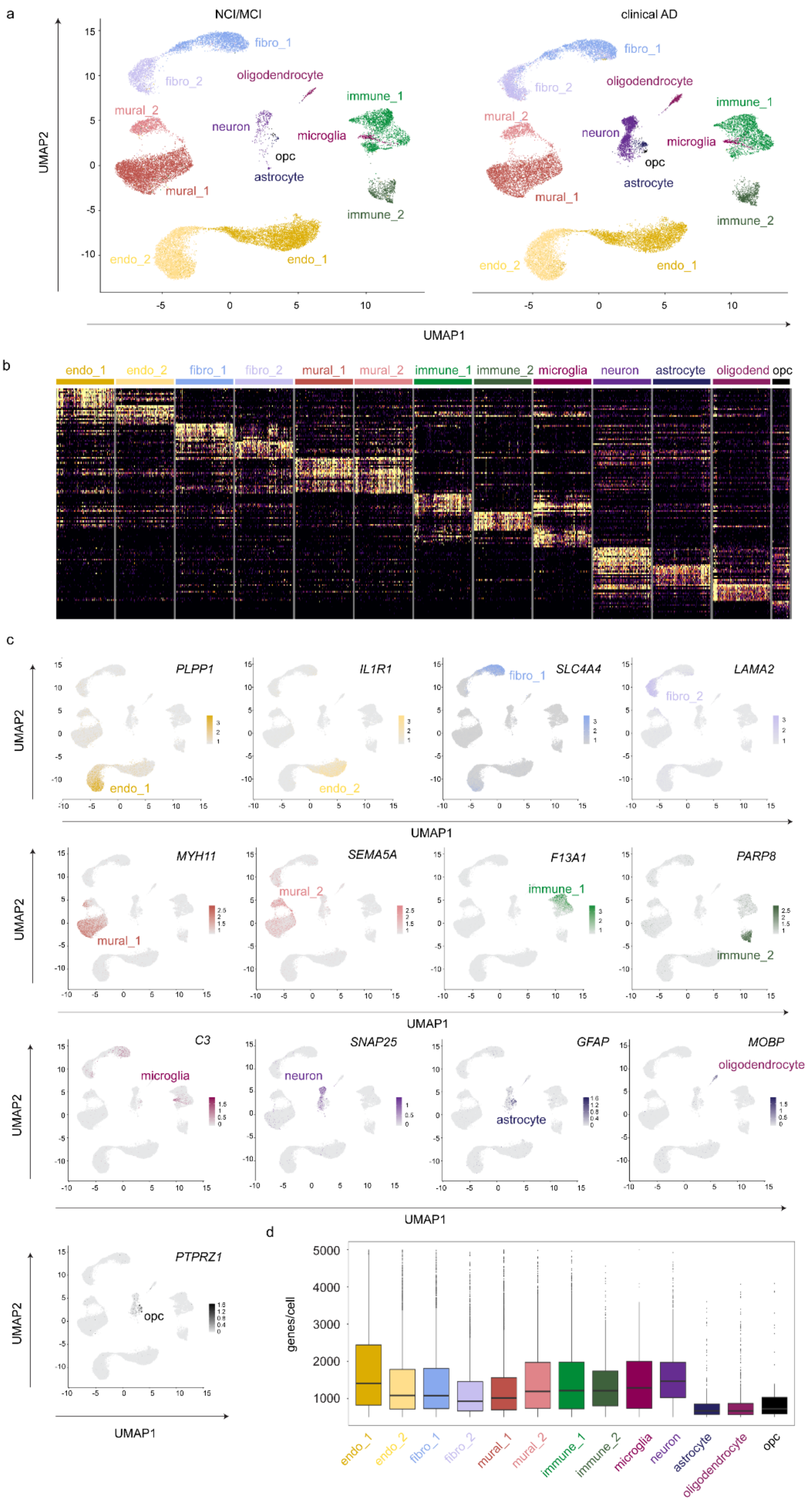


**Extended Data Figure 2. snRNA-seq reveals major cell types of the human leptomeninges and adjacent parenchyma.** (a) UMAP of meningeal and parenchymal cell types integrated across all donors, colored by coarse cell types, separated by NCI/MCI and AD. (b) Heatmap showing the top 10 markers per cell type across 100 randomly selected cells per cluster. (c) UMAP visualization of marker genes representing each cell cluster. (d) Boxplot shows the number of unique genes detected per nucleus in each cell type.

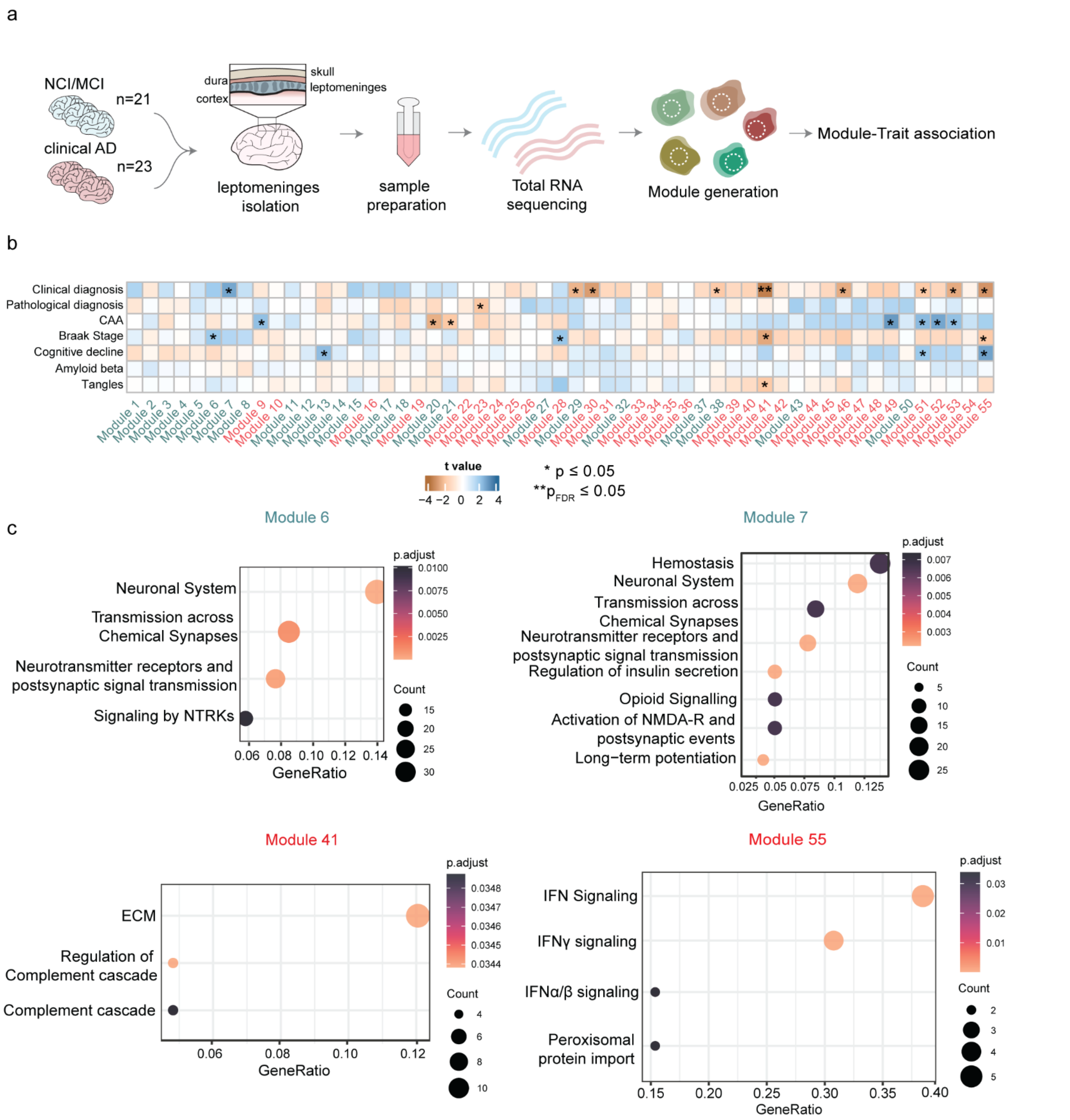


**Extended Data Figure 3. AD-associated gene modules**. (a) Schematic illustration of experimental design and module-trait association analysis. (b) Association between modules and AD-related traits, including clinical and pathological diagnosis, cerebral amyloid angiopathy (CAA), cognitive decline, plaques, and tangles. *; $p \leq 0.05$, **; $p_{FDR} \leq 0.05$ (c) Pathway analysis of the gene members of preserved modules 6 and 7, and non-preserved modules 41 and 55. Significant Reactome pathways are visualized ($p_{BH} < 0.05$).

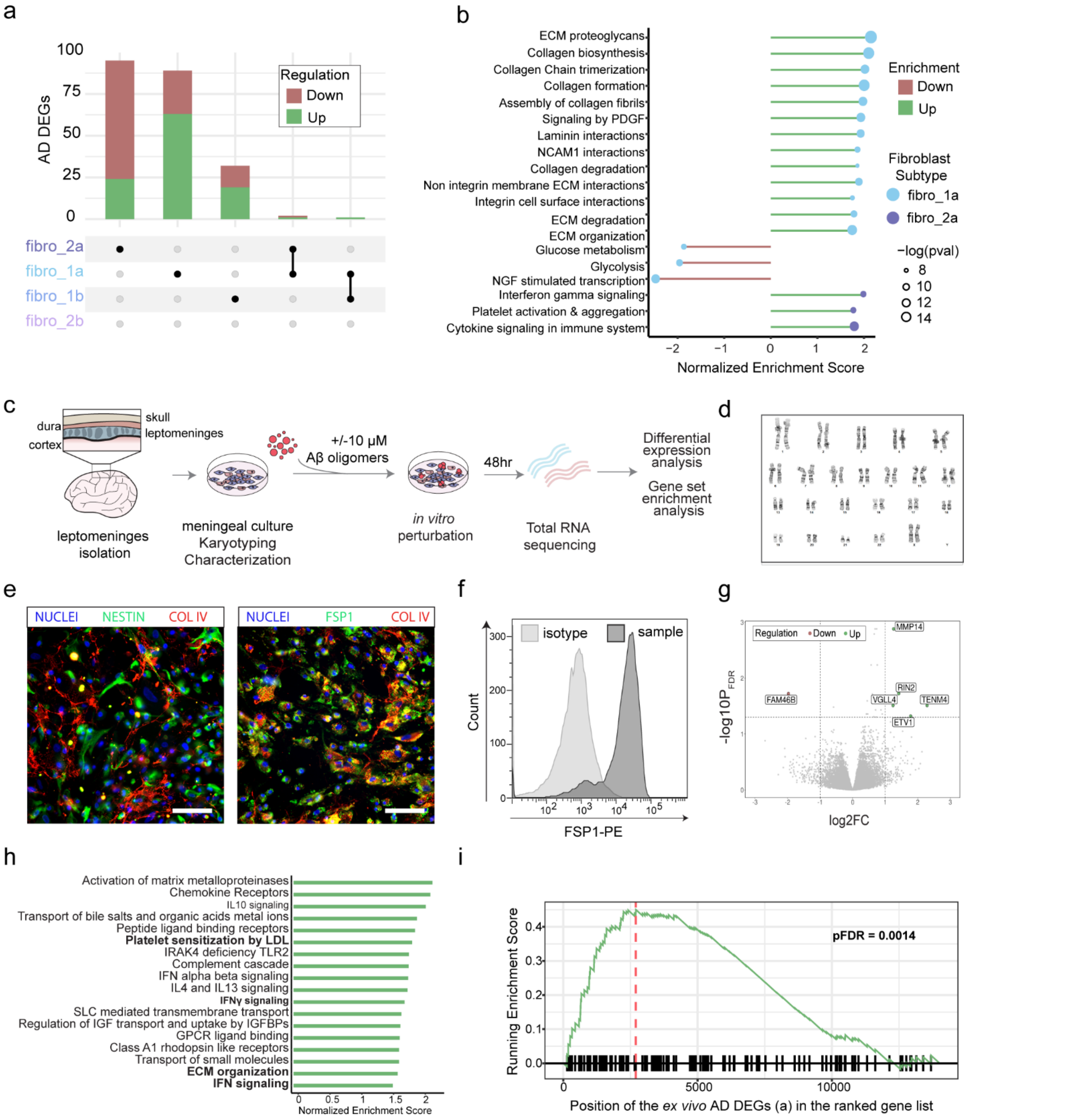


**Extended Data Figure 4. Aβ-treated leptomeningeal fibroblast-like cells show the AD gene signature of *ex vivo* fibroblasts.** (a) Differentially expressed genes (DEGs) between AD and control in each fibroblast subtype; threshold set at |logFC| > 0.3 and $p_{BON} < 0.01$. (b) Gene set enrichment analysis reveals significant terms ($P_{FDR} < 0.05$) for fibro_1a and 2a subtypes. (c) Schematic of leptomeningeal fibroblast line derivation and Aβ treatment paradigm. (d) Representative karyotype result of a derived fibroblast line. (e) Representative immunofluorescence images of cultured fibroblasts stained for nestin, collagen IV, and fibroblast-specific protein 1 (FSP1). Scale bar = 5μm. (f) Representative flow cytometry histogram showing the FSP1 expression of cultured fibroblasts. (g) DEGs in cultured fibroblasts upon Aβ treatment. (h) Gene set enrichment analysis of the DEGs showing significant terms ($P_{FDR} < 0.05$). (i) Enrichment score of the *ex vivo* fibroblast AD DEGs in the pre-ranked list of the *in vitro* Aβ-induced DEGs.

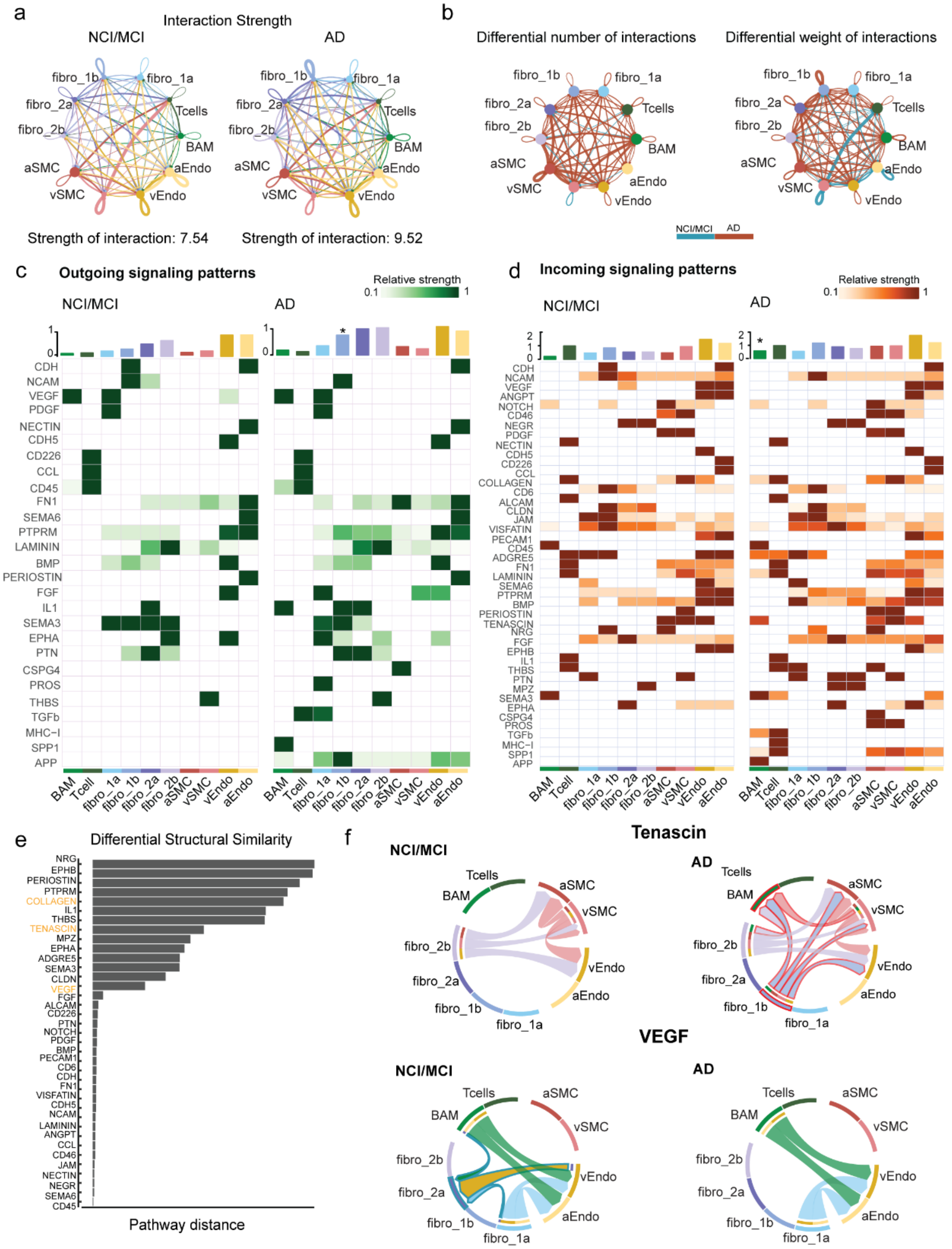


**Extended Data Figure 5. Altered intercellular communications in AD leptomeninges.** (a) Circular plots show the strength of inferred intercellular communications in control and AD groups. Autocrine (loops on top of each cell group) and paracrine (connections between cell groups) communications are displayed in each circular plot. (b) Differential number and weight of communications between the control and AD groups. Blue and red lines denote increased communications in control and AD, respectively. (c-d) Outgoing and incoming active signaling patterns across cell types in control and AD. The bar plots on the top show the sender or receiver's overall activities in each cell type, and the asterisks annotate significantly increased signaling in AD. Note that Fibro-1b and BAMs in AD display significantly increased outgoing and incoming activities, respectively, denoted by the black asterisk. Wilcoxon Rank-Sum $p < 0.05$ (e) Evaluation of similarity of signaling network topology of each shared pathway between the control and AD groups. (f) Circular plots show the altered Tenascin and VEGF pathways in AD. Differential interactions are highlighted in red for AD and blue for control.

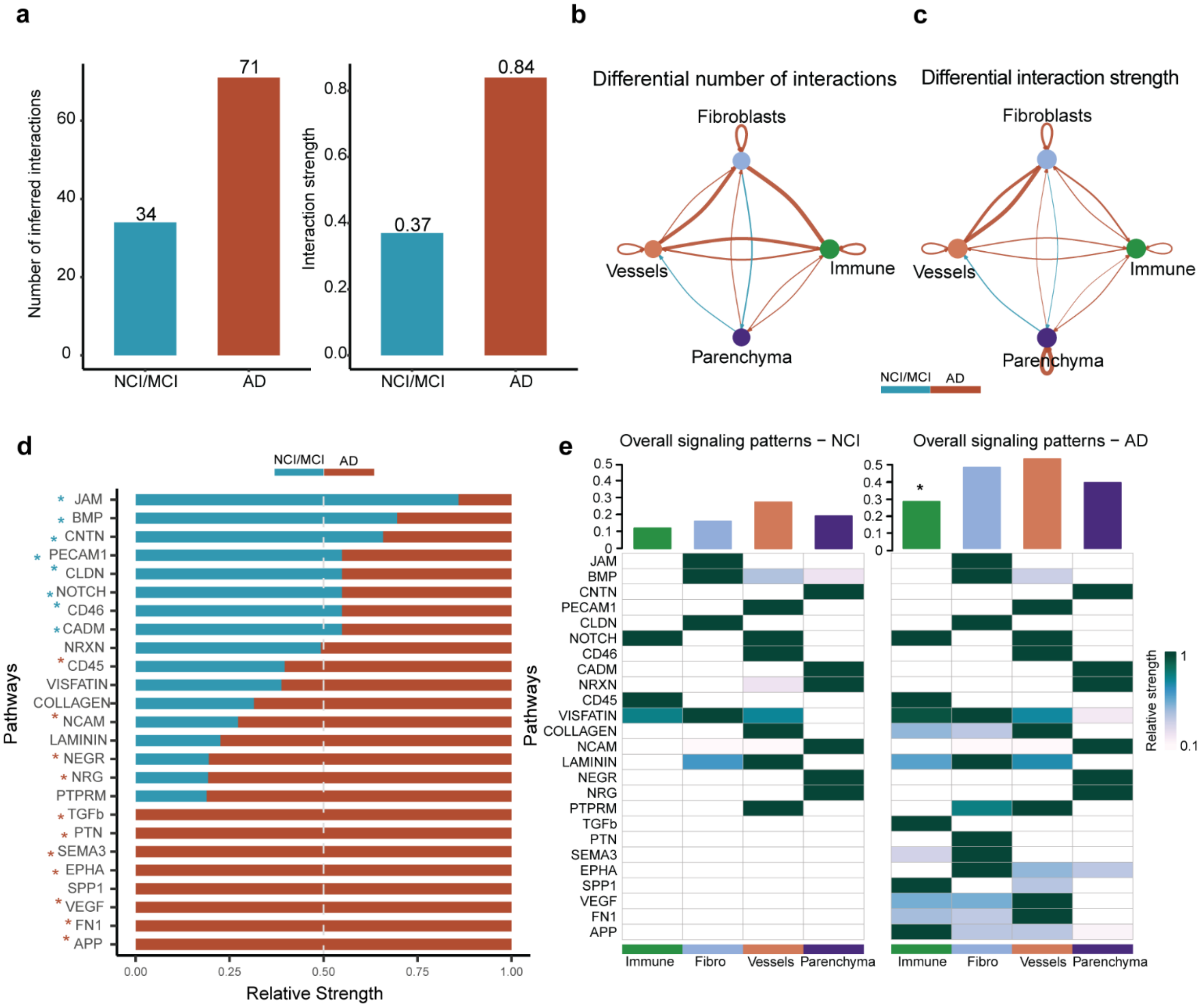


**Extended Data Figure 6. Altered intercellular communications at AD leptomeninges-parenchyma interface.** (a) Bar plots of the total number and strength of inferred interactions in control and AD. (b-c) Differential numbers and weight of interactions among parenchymal and leptomeningeal cells. Blue and red lines denote increased communications in control and AD, respectively. (d) The relative strength of 25 active signaling pathways at the leptomeninges-parenchyma interface in control and AD groups. Pathways annotated with blue and red stars are downregulated and upregulated in AD, respectively. Wilcoxon Rank-Sum $p < 0.05$. (e) The overall signaling patterns across cell types in NCI/MCI and AD. The bar plots on the top show the overall activity of each cell type. * denotes significantly increased activity.

**Supplementary Tables**

**Table S1**. Demographic characteristics of selected cases for *ex vivo* and *in vitro* experiments.

**Table S2**. Bulk RNA-seq Modules. **Related to Extended Figure 1**.

**Table S3**. Marker genes for major cell types. **Related to Figure 1 and Extended Figure 2**.

**Table S4**. DEGs for vascular trajectory analysis. **Related to Figures 2**.

**Table S5**. Marker genes for fibroblast subtypes. **Related to Figure 3**.

**Table S6**. Marker genes for T cell subtypes. **Related to Figures 4.**

**Table S7.** DEGs between microglia and BAMs. **Related to Figure 4**.

**Table S8**. Marker genes for BAM Clusters 1-3. **Related to Figure 5.**

**Table S9**. Selected GWAS genes and their hierarchical clustering. **Related to Figure 5.**

**Table S10.** Module-trait association. **Related to Extended Figure 3**.

**Table S11**. DEGs in AD by cell types. **Related to Extended Figures 3 and 4**.

**Table S12**. DEGs upon A-beta treatment. **Related to Extended Figure 4.**

**Table S13.** Primary antibodies and RNAscope probes.
